# Highly scalable and standardized organ-on-chip platform with TEER for biological barrier modeling

**DOI:** 10.1101/2023.08.02.551612

**Authors:** Hoang-Tuan Nguyen, Siiri-Liisa Rissanen, Mimosa Peltokangas, Tino Laakkonen, Jere Kettunen, Lara Barthod, Ragul Sivakumar, Anniina Palojärvi, Pauliina Junttila, Jussi Talvitie, Michele Bassis, Sarah L. Nickels, Jens C. Schwamborn, Sebastien Mosser, Prateek Singh

## Abstract

The development of new therapies is hampered by the lack of predictive, and patient-relevant in vitro models. Organ-on-chip (OOC) technologies can potentially recreate physiological features and hold great promise for tissue and disease modeling. However, the non-standardized design of these chips and perfusion control systems has been a barrier to quantitative high-throughput screening (HTS).

Here we present a scalable OOC microfluidic platform for applied kinetic in vitro assays (AKITA) that is applicable for high, medium, and low throughput. Its standard 96-well plate and 384-well plate layouts ensure compatibility with existing laboratory workflows and high-throughput data collection and analysis tools. The AKITA plate is optimized for the modeling of vascularized biological barriers, primarily the blood-brain barrier, skin, and lung, with precise flow control on a custom rocker. The integration of trans-epithelial electrical resistance (TEER) sensors allows rapid and repeated monitoring of barrier integrity over long time periods.

Together with automated confocal imaging and compound permeability testing analyses, we demonstrate the flexibility of the AKITA platform for establishing human-relevant models for preclinical drug and precision medicine’s efficacy, toxicity, and permeability under physiological conditions.

## 1. Introduction

Drug development is a lengthy and expensive process with a high failure rate, as only 10% of drugs entering clinical trials make it to the market ^1^. One of the main reasons for this failure is the poor predictivity of current preclinical models ^1–4^. To solve this major issue, a new generation of in vitro models has been developed, called organ on chip (OOC). Those allow the modeling of relevant human physiological features ^5–9^. By emulating perfusion, mechanical stimulation, and other physiological parameters essential to tissue and organ physiology, the OOC devices are expected to mimic physiological conditions better than conventional cell culture models.

OOC technology has advanced significantly in recent years, with numerous innovations that have improved these models’ functionality, versatility, and reliability. OOCs are microfluidic devices that aim to replicate the structure and function of human organs in a controlled environment, enabling researchers to study physiological processes and drug responses in vitro ^7,10^. One of the major advantages of OOCs is their ability to mimic the complex intercellular interactions that occur within organs, enabling researchers to study disease mechanisms and drug responses by targeting yet untested in vitro physiological parameters. For instance, a lung OOC may be used to assess chemotherapy response, or how air pollution affects lung function ^11^. Similarly, blood-brain barrier (BBB) OOCs can be used for PK/PD tests to assess the capacity of drug compounds to reach the brain or study the inflammation of brain vasculature and how it affects the onset of neurodegenerative diseases.

Current in vitro models partially recapitulate the in vivo microenvironment, but there is a strong body of literature indicating the physiologically relevant cues that guide tissue function. For instance, the flow has a significant impact on cells and tissue ^12^. It provides perfusion at lower flow rates, which helps to deliver nutrients and oxygen to cells, and removes waste products ^13–18^. At higher flow rates, flow induces wall shear stress (WSS) on cells and tissue, which can influence their function and behavior ^19–22^. Shear stress is involved in various physiological processes such as blood vessel dilation ^23^, inflammation ^24^, and wound healing ^25^. Additionally, WSS has been shown to affect cell differentiation ^26^, proliferation ^27^, and gene expression, which can have implications for tissue engineering and disease modeling. Furthermore, maintaining near-physiological concentrations of autocrine and paracrine signals requires a low medium-to-cell ratio within in vitro culture systems.

OOC technology has the potential to aid the development of new therapies, however, their development has been carried out at the expense of throughput ^28,29^. Indeed they require more specialized and labor-intensive procedures to fabricate and maintain. Furthermore, OOCs are rarely compatible with the industry-standard workflow and infrastructure because they require different equipment, protocols, and expertise compared to traditional cell culture models. Those adoption barriers decrease the likelihood for pharmaceutical companies to adopt and integrate OOC technology widely into their drug development pipelines.

To address those limitations, we present a scalable - from high to low throughput – automation-compatible OOC platform for applied kinetic in vitro assays (AKITA). It is suited for the modeling of endothelial-epithelial barriers, and the culture of organoid or cell migration. Its format is compatible with ANSI/SLAS standard 96-well (24 units of OOC) and 384-well (128 units of OOC) plates, compatible with common laboratory workflows. For high ease of manipulation, the flow is induced by a tubeless system - a rocker - whose angle can be set up to precisely control the flow rates. Furthermore, full plate impedance spectroscopy allows the real-time recording of barrier integrity via trans-epithelial electrical measurements (TEER).

As an example, we show the generation of skin-, BBB-, and lung-on-chip were generated and validated by a fluorescent permeability assay, TEER measurements, and automatized confocal microscopy. This work aims at providing seamless integration of OOC technology into the current disease modeling and drug study pipeline of both academic research and the pharmaceutical industry.

## 2. Materials and methods

### 2.1. Microfluidic plate design

The AKITA plates comply with the ANSI/SLAS cell culture plate standard, having the form and dimensions of the 96 or 384-well plate. The plate incorporated 24 (96-well plate) or 128 (384-well plate) microfluidic units (one unit corresponds to one test or experimental condition). Each unit regroups three wells to form the complex structures underneath. In this paper, we mainly explain the design and manufacturing of the AKITA Plate 96. For each unit (three linked wells) the middle well plays as an open-top chamber; the other two wells are linked together through a microchannel on the bottom. At the area where the microchannel is crossing the bottom of the middle well, a semi-permeable polyethylene terephthalate (PET) membrane is integrated to separate the two compartments. The width and height of the microchannel were designed to gradually ramp up from 0.3 mm to 6 mm and from 0.15 mm to 0.3 mm from the 2 inlets toward the middle area, to ensure the smooth liquid laminar flow and avoid any turbulence. The membrane was 9 μm thick with a pore size of 3 μm, acting as an interfacing layer between the microchannel and the open-top chamber. The design of the AKITA Plate allows a bilateral flow through the microchannel using a rocker platform to pass the liquid from one well to the other while keeping the open-top chamber static with no liquid circulation. This combination of both flow (microchannel) and static (middle well) conditions acting on both sides of the membrane give a versatile platform for multiple in vitro co-culture models where one cell type can experience the flow-induced shear stress while the other cell type remains static or exposed to very low inertial flow.

An additive manufacturing-assisted soft-lithography procedure was developed to fabricate the complex structure of the plate. With the membrane clamped in between, the 3D printed 2-part negative mold was cast with a 10:1 mixture of the Polydimethylsiloxane (PDMS) prepolymer and curing agent. Finally, the demolded PDMS structure was bonded onto a 100 μm thick PDMS film using a plasma cleaner (Harrick Plasma Inc., US). The process yields a full plate with multiple layers while reducing the need for many bonding processes between layers.

### 2.2. Label-free TEER assessment of barrier integrity

To monitor the development of the tissue barriers formed on the AKITA plate 96, we used the AKITA Lid device developed by Finnadvance, to enable high throughput, label-free, non-invasive TEER measurements of 24 channels simultaneously. The AKITA Lid consisted of an analog front-end for an impedance analyzer with a frequency sweeping spectrum from 100 Hz to 200 kHz, a digital back-end for main function control, and a detachable electrode array. The electrode array circuit facilitated 144 gold-plated chopstick-like electrodes. The tetrapolar configuration was used for the sensing, there are one pair of excitation electrodes and one pair of measurement electrodes. Theoretically, the electrodes of each pair are placed on the opposite side of each other, i.e., on both sides of the membrane, via the inlet/outlet reservoirs and open-top chamber. The impedance values are measured and transmitted to the computer software in three ways, depending on the laboratory’s accessibility: Wi-Fi, serial port, and micro-sd card. The measurements were performed before the medium change, at 37 °C, to reduce the variations in temperature. For the cells under air-liquid interface (ALI), the culture medium was added to the open-top chamber just for measurement and removed after it. TEER values were calculated by fitting the recorded impedance data to an electrical equivalent circuit of tissue barriers described elsewhere ^30^. In short, the circuit includes a serial combination of a resistor representing culture medium resistance, a resistor in parallel with a capacitor representing the cell layer, and a constant phase element CPE (electrode-medium interface).

### 2.3. Computational studies

#### 1) Gravity-driven fluid dynamic simulation

Flow rate, shear stress, and flow streams were simulated using finite element analysis (COMSOL Multiphysics). A 3D computer-aided design (CAD) model of the chip was created from Solidworks (Dassault Systèmes) and imported into COMSOL. Free and porous medium flow physics was selected for modeling the fluid flow which is governed by Navier Stokes and Brinkman equations ^31^. The incompressible flow was used with Phosphate buffer solution (dynamic viscosity = 0.001 Pa.s, Density = 1005.6 Kg/m3) as the fluid material. PDMS was selected as the boundary walls and a no-slip condition was applied to the walls. Porous material was included, and the membrane was selected as a porous medium. The gravitational force was included as a volumetric force which induces hydrostatic pressure for driving fluid inside the channel.

The moving mesh was applied for simulating the rocker motion where the hydrostatic pressure changes with respect to the tilting angles. A normal mesh was used, and a time-dependent study was implemented for 150 seconds to study the velocity, flow rate, and shear stress from 10° to 30° tilting angles. Shear stress was calculated by multiplying the shear rate and fluid viscosity whereas the flow rates were determined by taking the surface integral of fluid velocity in the microchannel under the membrane.

#### 2) Electrical simulation for TEER sensitivity

Sensitivity field distribution on the membrane where the cells grow was simulated using COMSOL Multiphysics. The 3D geometry of the chip with submerged electrodes was designed in Solidworks and imported into COMSOL. Electric field physics was used for studying sensitivity on the cell layer at different electrode setups (tetrapolar configuration with 4 or 6 electrodes). The electrodes are 0.4 mm in diameter, and 10 mm in length, with gold as the material. Alternative current (AC) of 10 µA was injected through the Current Carrying (CC) electrodes and Voltage Sensing (VS) electrodes were set as floating potential. The model was simulated at TEER values ranging from 10^1^ to 10^3^ Ω.cm^2^, for covering the TEER range of the most popular in vitro tissue barrier models. A wide-range frequency domain (10E1 – 10E6 Hz) study was used. Parametric sweeps at different TEER values were simulated and compared with the help of sensitivity distribution fields (at different electrode diameters). Fine mesh was used in solving all the models.

### 2.4. Cell culture

Primary mouse Brain Microvascular Endothelial Cells (mPBMECs, CellBiologics) were cultured in a complete Endothelial Cell Medium (CellBiologics) in petri-dish pre-coated with 0,2% gelatin (Merck Life Science). Immortalized human cerebral microvascular endothelial cell line (hCMEC/D3 Merck Life Science) were cultured on 250µg/ml collagen-I (Advanced Biomatrix) coated petri dish in EndoGRO-MV complete culture medium (Merck Life Science) supplemented with 1% PenStrep (Gibco) and 1ng/ml basic fibroblast growth factor (Merck Life Science). Adenocarcinoma human alveolar basal epithelial A549 cells (Sigma) were cultured in Ham’s F-12 K medium (Fisher Scientific) supplemented with 10 % Fetal bovine serum (FBS) (Serana), 1 % penicillin/streptomycin (Gibco), and 2 mM L-Glutamine (Sigma). Human umbilical vein endothelial cells (HUVECs) were kindly provided by prof. Eklund from Oulu University. HUVECs were cultured on an attachment factor (Gibco) coated petri dish in Endothelial cell growth medium 2 (ECGM2) with a supplement kit (Promocell), 10% FBS, and 1% PenStrep. Normal human dermal adult fibroblasts (NHDFa, Lonza) were grown in DMEM (Gibco) supplemented with 10% FBS and 1% PenStrep. Normal human epidermal neonatal keratinocytes (NHEKneo, Lonza) were cultured on KGM Gold keratinocyte growth medium (Lonza). In all the experiments, cells were cultured at 37 °C with a 5 % CO2 concentration.

### 2.5. Blood-brain barrier-on-chip

Mouse blood-brain barrier (BBB) model was established with a tri-culture of mouse brain microvascular endothelial cells cultured inside the microchannel in a complete endothelial cell medium, and a mixture of primary mouse brain astrocytes (mAs, ScienCell) and mouse primary brain pericytes (mPe, iXCells) cultured on the open-top chamber in 1:1 mixture of Astrocyte Medium-animal (ScienCell) and pericyte growth medium (iXCells). Whereas the human BBB model was established with hCMEC/D3 cultured inside the microchannel with EndoGRO-MV complete culture medium. The triculture model of the human BBB was established with a mixture of human primary cerebellar astrocytes (hAS, ScienCell) and human primary brain vascular pericytes (hPe, ScienCell) cultured on the open-top chambers in 1:1 mixture of complete astrocyte medium (ScienCell) and complete pericyte medium (ScienCell).

AKITA Plate microchannels and open-top chambers were coated with 70µL and 50µL respectively, of 100µg/ml human fibronectin (Corning) diluted in phosphate-buffered saline (PBS, Gibco). The coating solution was incubated for 4 hours at +37 °C, 5% CO_2_, following an overnight incubation at +4 °C. On the day of the cell loading the coating solution was removed and replaced with the suited culture medium. The endothelial cells in both human and mouse models were seeded to microchannels in 2E6 cells/ml cell concentration. The plate was flipped upside down and incubated upside down for 4 hours at +37 °C, 5% CO_2_, allowing the cells to attach to the top side of the microchannel. Astrocytes and pericytes in both models were seeded to the open-top chambers in 1E6 cells/ml concentration to form a triculture model. All cells for both human and mouse BBB were seeded to the AKITA Plate straight after thawing from liquid nitrogen.

The plates were either cultured in static conditions or on the AKITA Wave rocker with a 20-degree tilting angle, 1 RPM speed, and 40-second hold time at extreme angles. The medium was replenished every day in the microchannels and open-top chambers. The cells were grown on AKITA® Plate for 5-7 days prior to permeability assay or fixing.

### 2.6. Blood-Brain barrier and midbrain organoid co-culture

The neurovascular unit was established on the AKITA Plate 96 with a coculture of hCMEC/D3 cells inside the microchannel and a midbrain organoid (OrganoTherapeutics) in the open-top chamber. Briefly, the endothelial barrier with monoculture of hCMEC/D3 cells was established as described at human BBB formation above, only using hCMEC/D3 cells. The human BBB was cultured for seven days under flow conditions allowing a barrier formation. On day seven of the experiment the medium on the open-top chamber was changed to midbrain organoid culture medium (OrganoTherapeutics) and the midbrain organoids were loaded to the open-top chambers one by one. To assess if the viability of the organoids or the simple BBB remains stable during the coculture period, midbrain organoids and BBB were also monocultured on the AKITA Plate as a control. The formed neurovascular unit was further cultured for 48h.

### 2.7. Skin-on-chip

The skin-on-chip model was established by culturing human dermal fibroblast in a collagen mixture on top of the porous membrane, following human epidermal keratinocyte culture on top of the dermal layer. The AKITA plate units were coated with 100 µg/ml collagen-I solution diluted in PBS as explained in the blood-brain barrier model. On the day of cell seeding, the fibroblasts were diluted to a final concentration of 1E6 cells/ml in a mixture with 100 µg/ml of collagen-1 and culture medium and loaded into the open-top chambers. After 3 days of culture, keratinocytes were added on top of the fibroblast layer in a concentration of 1E6 cells/ml. The cells were left to attach in static conditions in the incubator overnight. Monoculture units were also cultured as a control.

After attachment, the plates were either cultured in static conditions or on the AKITA Wave rocker (20° tilting angle, 1 RPM speed, and 40-second hold time) to achieve dynamic flow. After 3 days of seeding keratinocytes, the units were exposed to the air, and an air-liquid interface (ALI) was created. During the coculture, the mixture of 1:1 DMEM and KGM medium was used for the model, and the medium was changed daily. The model was cultured for 11 days under ALI, following the permeability assay, and fixing.

### 2.8. Lung-on-chip

The lung-on-chip model was established with a coculture of HUVECs cultured inside the microchannel and A549 cultured on the open-top chambers. The AKITA plate’s units were coated with 100 µg/ml collagen-I diluted in PBS as explained in the blood-brain barrier model. Then A549 cells were seeded to the open-top chambers in 0,25E6 cells/ml concentration. The plate was incubated statically for 4 hours at +37 °C, 5% CO_2_, allowing the cells to attach. Next, HUVECs were seeded to microchannels in 2E6 cells/ml cell concentration, and the plate was flipped upside down and incubated overnight at +37 °C, 5% CO_2_. On the next day, the plate was flipped back right side up.

As with the BBB and skin models, the plates were either cultured on static conditions or on the AKITA Wave rocker with the 20° tilting angle, 1 RPM speed, and 40-second hold time at extreme angles. The medium was replenished every day in the microchannels and open-top chambers. After culturing cells for 5 days under the submerged environment with culture medium on both sides of the membrane the medium was removed from open-top chambers, and A549 epithelial cells were exposed to the air. As a comparison, control units were cultured under a submerged environment during the whole experiment. The culturing was continued for an additional 4 days for full barrier maturation.

### 2.9. Permeability assay

Fluorescent-labeled molecules, 0,4 kDa sodium fluorescein (Sigma), 70 kDa TRITC-dextran (TdB Labs), and 4 kDa FITC-dextran (TdB Labs) were used to test the barrier permeability of all the organ models. Briefly, the medium of the open-top chamber was first changed to 70 µl of fresh medium, and medium background samples were collected. Next, reservoirs of the microchannel were emptied and 100 µl of tracer solution in culture medium (0,5 mg/ml for dextran, 0,05 mg/ml for sodium salt) was added to the reservoirs. The AKITA Plate was placed on the rocker and incubated for either 60 or 120 min. After incubation, samples from both the open-top chamber and microchannel reservoir were collected to a microtiter plate.

The fluorescent intensity of the samples was measured with a plate reader (TECAN infinite M1000Pro). For 4 kDa and 0,4 kDa tracers, 498 nm excitation wavelength, and 518 nm emission wavelength were used. For the 70 kDa tracer 554 nm excitation wavelength and 578 nm emission wavelength were used. The permeability coefficient at desired time point was determined as previously reported ^32^.

Briefly, apparent permeability (Papp) was calculated using the following equation (1):

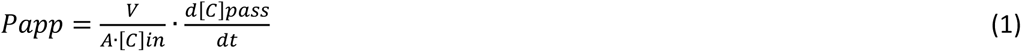

Where *V* is chamber volume of the passed tracer (cm^3^), *A* is membrane surface area (cm^2^), *[C]in* is initial tracer concentration (ng/ml), *d[C]pass* is passed tracer concentration (ng/ml), and *dt* is the time of the experiment (s).

### 2.10. Reversible barrier opening

For reversible barrier opening assay, a BBB model was established as described in mouse blood-brain barrier-on-chip formation with few exceptions, instead of fibronectin, a 250µg/ml collagen-I was used as coating solution and the barrier was established without pericytes. On day seven of the experiment, first the initial permeability of the established BBB was assessed with a permeability assay with a 70kDa dextran as described above. After the permeability assay, the normal culture medium was changed to a hypertonic culture medium (Finnadvance) and incubated for 1 h. The opened barrier permeability was measured with 70kDa dextran solution supplemented with a hypertonic culture medium. After this, the culture medium was changed yet again to the normal isotonic medium to allow the cells to recover for 1 hour from the opening of the barrier, to ensure the recovery the permeability assay was once again performed with the 70kDa dextran.

### 2.11. Live staining and imaging

During the experiments, cell confluency and viability were observed with live staining and imaging. For this purpose, Calcein-AM (Fisher Scientific) was used. Calcein-AM was diluted into a fresh culture medium with a final concentration of 2 µM and the culture medium on both microchannels and open-top chambers was replaced with a staining solution, following 30 min incubation. Next, the fresh culture medium was changed, and the units were imaged with a fluorescence microscope.

### 2.12. Immunofluorescence staining

At the end of the experiments, the organ-on-chip units were fixed with 4% PFA and continued to the immunofluorescence staining. To minimize nonspecific binding the samples were blocked for 4 h at room temperature with a blocking buffer containing 1% of Tween20 (Fisher Scientific) and 3 % of BSA (VWR) in PBS. For skin-on-chip, permeabilization with 0.1% Triton X100 (Fisher scientific) in PBS was performed for 10 minutes prior to blocking.

Primary antibodies used, listed in Table 1, were diluted into a blocking buffer, and incubated overnight at +4 °C. After incubation, the cells were washed three times with PBST (1% Tween20 in PBS). Similarly, secondary antibody dilutions were prepared in a blocking buffer and incubated overnight at +4 °C, stained cells were washed 3 times with PBST. Membranes of BBB and lung models with stained cells were cut out of the AKITA Plate and mounted with Immu-Mount (Fisher Scientific) between microscopy glass slides for imaging. Skin-on-chip units were imaged directly on the Plate.

**Table 1:** immunofluorescence staining reagents.

|  | Primary antibodies | Dilution | Secondary antibodies | Dilution |
| --- | --- | --- | --- | --- |
| Lung | Phalloidin CF660R (BioNordika) | 1:200 | - | - |
|  | MUC5AC, anti-human, Mouse mAb(Fisher Scientific) | 1:100 | CF488A Goat Anti Mouse IgG (H+L), Highly Cross-Adsorbed (min X Rat) (Biotium) | 1:2000 |
|  | - | - | DAPI (Biotium) | 1:1000 |
| Blood-brain barrier | Phalloidin CF594 (Biotium) | 1:200 | - | - |
|  | CD31 ((PECAM-1), anti-mouse, Goat pAb (Bio-Techne) | 10µg/ml | CF488A Goat Anti-Mouse IgG (H+L), Highly Cross-Adsorbed (min X Rat) (Biotium) | 1:2000 |
|  | - | - | DAPI (Biotium) | 1:1000 |
| Skin | Phalloidin CF594 (Biotium) | 1:200 | - | - |
|  | Collagen IV, anti-human, Rabbit pAb (Abcam) | 1:200 | CF680 Donkey Anti-Rabbit, Highly Cross-Adsorbed (Biotium) | 1:2000 |
|  | Loricrin, anti-human/anti-mouse, Rabbit pAb (Abcam) | 1:200 | CF680 Donkey Anti-Rabbit, Highly Cross-Adsorbed (Biotium) | 1:2000 |
|  | - | - | DAPI (Biotium) | 1:1000 |

### 2.13. Microscopy and image analysis

For live imaging, either Olympus CellSense time-lapse imaging system with an Olympus XM10 CCD camera and 4x objective with a GFP filter, or Zeiss Axio Zoom.V16 fluorescence stereo microscope with Hamamatsu Orca Fusion BT (monochrome) and Zeiss Axiocam 305 (color) cameras with PlanApo Z 0.5x/0.125 objective were used. For fixed BBB and lung samples, imaging was performed using a Leica TCS SP8 confocal with a DMI8 microscope using LAS X 3.5.2 acquisition software. The used objective was HC PL APO 20x/0.75 CS2 DRY with excitation of 405 nm, 489 nm, and 661 nm solid-state lasers. Fixed skin-on-chip units were imaged with Leica STELLARIS 8 DIVE multiphoton confocal microscope using a Spectral fluorescence detector (3 HyD S). The used objective was HC PL APO 20x/0.75 CS2 (Air) with excitation of 405 nm, 561 nm, and 638 nm lasers.

Scanning electron microscopy (SEM) and Transmission electron microscopy (TEM) samples were prepared and imaged in collaboration with the electron microscopy core facility of Biocenter Oulu. Cells were fixed with fixative (1 % glutaraldehyde, 4 % formaldehyde in 0.1M phosphate buffer, provided by Biocentre) and stored at +4 °C before sample processing. For SEM imaging, Sigma HD VP FE-SEM was used with 5 kV electron high tension and a pixel size of 44,66 nm. For TEM imaging, Tecnai G2 Spirit 120 kV TEM with Veleta and Quemesa CCD cameras were used.

Image analysis was made with ImageJ (Rasband., 2018) and Bitplane Imaris 3D software (Imaris., 2022) for fluorescence and confocal microscopy images, respectively. All the imaging was performed with the collaboration of Biocenter from the University of Oulu. Construction of the schemes was performed using Biorender software.

### 2.14. Statistical analysis

All data are presented as mean ± standard deviation. For all experiments, “N” represents the experimental repetitions. The homoscedasticity and homogeneity of variances were tested with the Shapiro-Wilk test and Bartlett test from all the collected data. Two-way analysis of variance (ANOVA) and Tukey’s multiple comparisons test were used to test differences between conditions. For inhomogeneous data, the Kruskal-Wallis test and the Dunn test with Bonferroni correction were used. The confidence levels of p<0,05 (*), p<0,01 (**), and p<0,001 (***) were used as significant differences in all statistical tests. R 4.2.3 software (R Core Team, 2023) was used for data analysis. The exact number of repeats performed for each experiment is indicated in the corresponding figure legends.

## 3. Results and Discussion

### 3.1. Microfluidic plate design and manufacturing

The AKITA Plate 96 comprises three main layers, the top reservoirs, the track-etched PET membrane, and the bottom microchannel (Fig. 1a-c). Due to the complexity of multiple layers, non-planar structures, and material characteristic differences, the fabrication of the plate was facing many challenges. However, we have optimized the production process to create a monolithic plate with an integrated membrane, without the existence of layer gaps, and eliminate the bonding steps between layers. As a result, a standardized well plate format OOC platform was designed and manufactured to enable a wide range of physiological barrier models in vitro (Fig. 1d).

**Figure 1.**
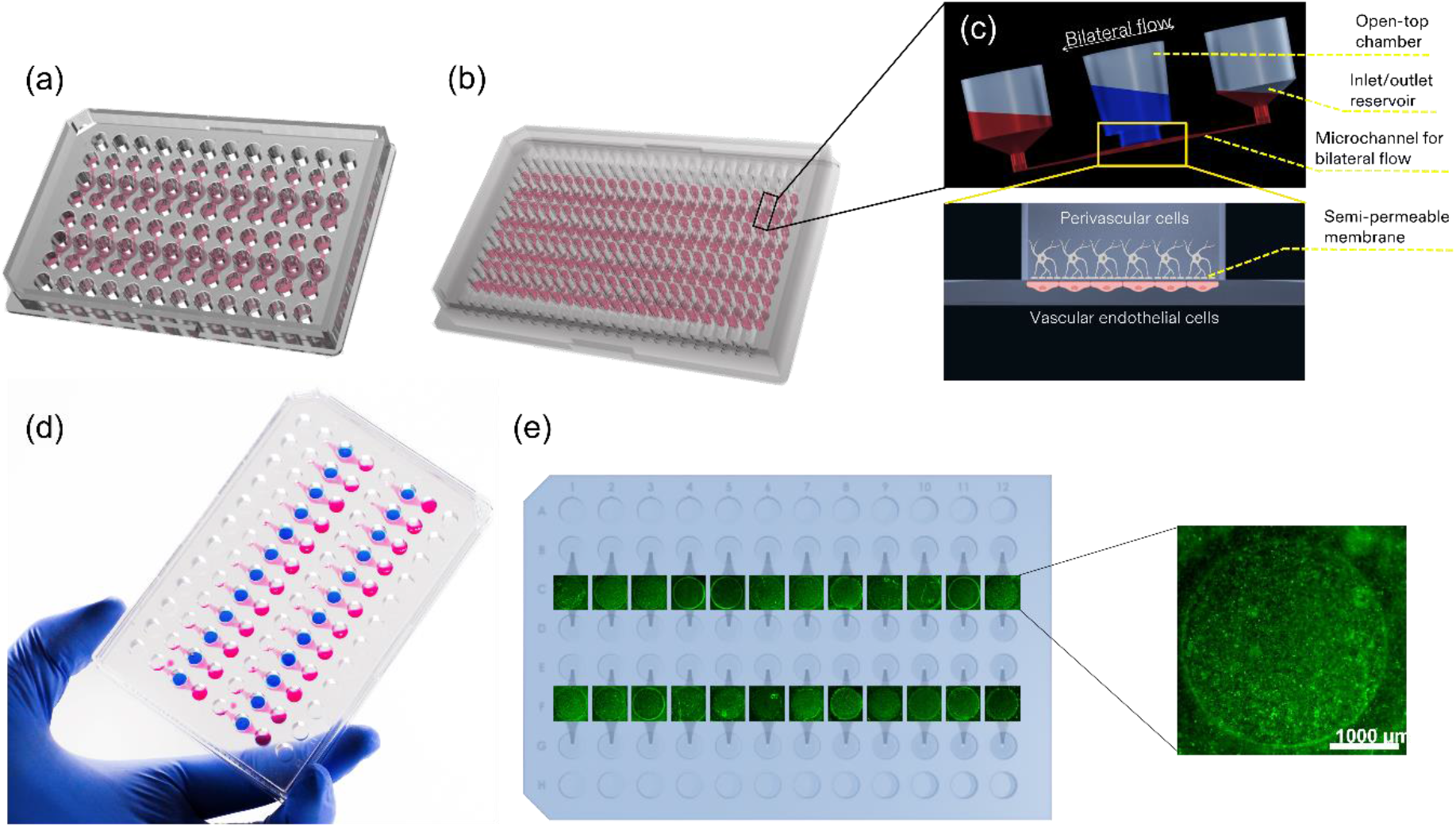
AKITA microfluidic plate design. (a) 3D rendered AKITA Plate 96 with 24 culture units. (b) 3D rendered AKITA Plate 384 with 128 culture units. (c) Side view of a culture unit on the AKITA Plate. The open-top chamber provides static culture conditions while the microchannel enables dynamic culture with the medium recirculated between the two inlet/outlet reservoirs. The semi-permeable PET membrane allows the cells on the open-top chamber and the microchannels to interact and form physiological barriers. (d) The manufactured AKITA Plate 96 is loaded with red dye in the microchannels and blue dye in the open-top chambers. (e) Top view of the AKITA Plate 96 matching with 24 live CAM staining images. The CAM signal shows the cells living in the membrane regions.

Figures 1a & 1b show the 3D rendered full plate designs of the AKITA Plate 96 (24 individual microfluidic units) and AKITA Plate 384 (128 individual microfluidic units). Within the context of this paper, we only discuss the results from the AKITA plate 96. Figure 1c illustrates how multiple layers of cells/tissues are interfacing with each other through the thin semi-porous membrane while coupling both the dynamic flow and static conditions for adapting to different cell types. With 24 (co)culture units on one plate, the experiment throughput was scaled up significantly compared to normal microscope slide-sized OOC devices.

The easy-access design with inlet/outlet reservoirs, and open-top chambers that are compatible with multichannel pipette or robotic liquid handling systems made the AKITA Plate 96 a suitable platform for high throughput screening applications. Microscopy imaging is an essential readout method for every cell-based model, Figure 1e shows an array of 24 live images of the cells’ Calcein AM signal at 4X objective taken from the AKITA Plate. While current 3D organ-on-chip models are difficult to be imaged due to their complex, non-standard 3D structures, our plate design enables high-resolution fluorescent or confocal fluorescent imaging thanks to the consistent cell culture constructs on the membrane across the units on the same plate and between multiple plates. Furthermore, the short focal distance (400 μm) from the plate substrate to the membrane where the cells are growing also plays an important role in the adaptation to most of the long working distance objectives of existing microscopy systems.

### 3.2. Pumpless flow optimization

Flow control is an important factor to establish the cell culture on-chip. Here we aim to eliminate the use of pumps for our high throughput platform as pump-based systems require a complex setup involving tubing, connectors, and valves, which can be challenging to assemble, maintain and scale up. With the AKITA Plates, we used a simpler approach to induce the flow through the microfluidic channels of the plate through the periodic tilting of the AKITA Wave rocker. Due to the hydrostatic pressure differences between the two reservoirs during the tilting periods, the liquid flows from one inlet to the other, bi-directionally. We performed the flow optimization by both in silico simulation and practical experiment (Fig. 2h) to ensure the precise flow profiles for specific tissue models. Figure 2a illustrates the consistent flow streams at the free-standing membrane region where the barriers are formed. As a result, a uniform shear stress distribution was observed, thus ensuring the same responses from the cells across the membrane area (Fig. 2b).

**Figure 2.**
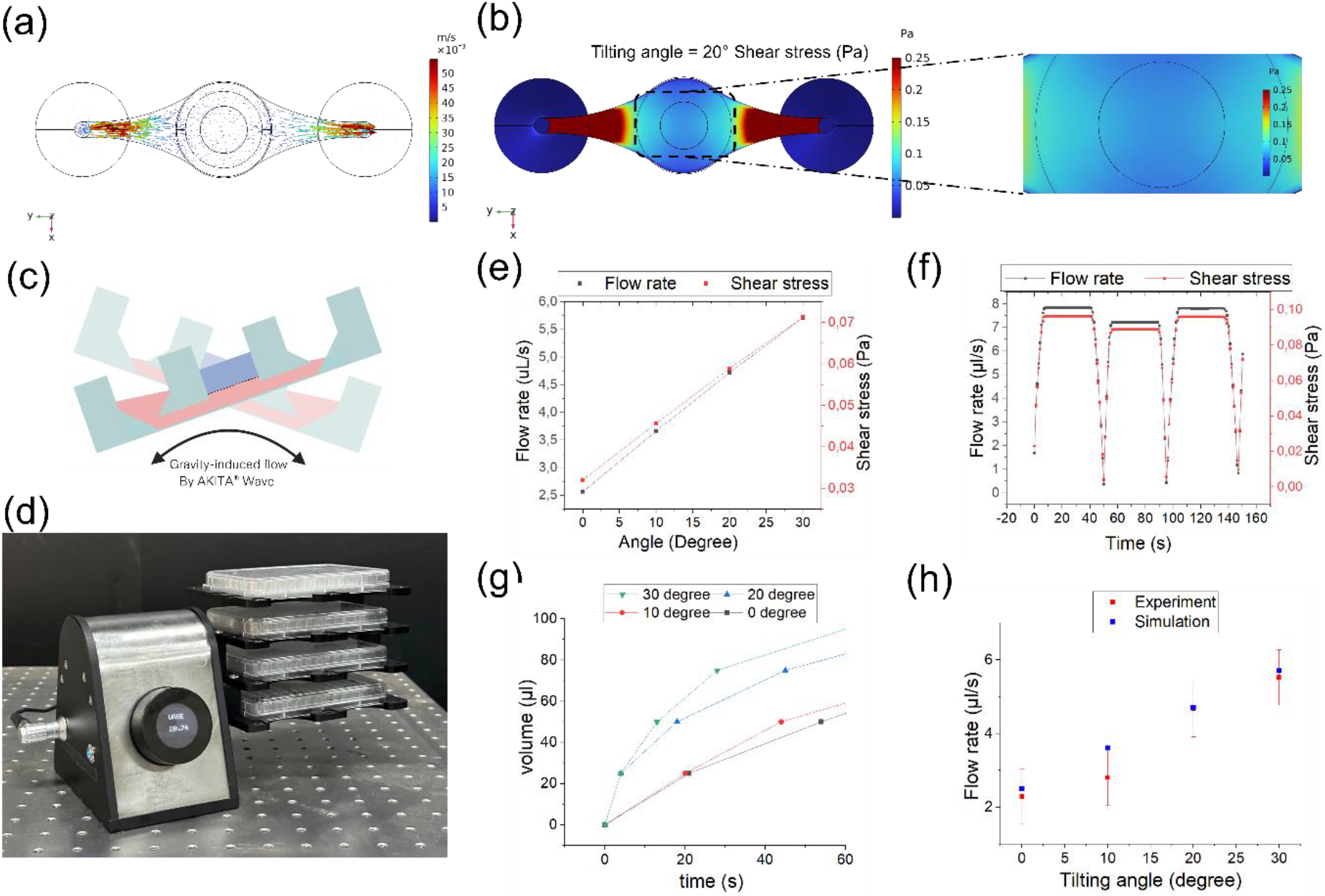
Bilateral flow simulations. (a) Arrow volume represents the velocity field of fluid inside the microchannel at the 20° rotation angle. (b) Surface plot showing the shear stress at the ceiling surface of the microchannel at the 20° tilting angle. Shear stress under the membrane area (zoomed-in picture) is uniform but smaller compared to the shear stress at the narrow regions of the microchannel. (c) Tilting motions of the plate due to the rocker’s movements. (d) The AKITA Wave rocker with 4 AKITA Plates stacking for dynamic cell culture. The rocker’s trays are designed to be compatible with ANSI/SLAS microtiter plates and robotic gripping arms. (e) Calculated values of the average shear stress at the membrane region and flow rates at different tilting angles. (f) Flow rate and average shear stress on the membrane over time during the rocker’s working cycles (tilting angle = 20°, rotation speed = 1 RPM, holding time on the left and right angles = 40 seconds). (g) The volume of liquid turnover at different tilting angles. (h) Comparison between simulation and experimental values of the microchannel’s flow rate.

As shown in Figure 2c, the plate is tilted from left to right periodically, yielding the changes in the height of the medium reservoir, increasing the volume of liquid passing through the channel over time (Fig. 2g). Furthermore, as the tilting angle is raised, the shear stress is linearly increased. Figures 2e & 2f show the linearity between the flow rate and shear stress, at different tilting angles and over time. Especially, figure 2f illustrates the steady flow rate periods when the rocker is held at a certain angle for 40 seconds. Based on those computed values, with different tissue types, the tilting angles, tilting time, and duty cycles can be optimized to give suitable shear stress and flow rate for the cells to grow. For instance, with our chip design, a rocker profile consisting of a tilting angle of 20°, 1 RPM speed, and 40 seconds holding time at the extreme left and right positions would yield 70 uL of medium passing through the microchannel in each period, and an average shear stress of 0.06 Pa across the membrane area.

### 3.3. Optimization of the electrode setups for TEER measurements and comparison with standard equipment

To implement the TEER measurements for the AKITA Plate 96 we used the AKITA Lid device, developed by Finnadvance Ltd. The AKITA Lid consists of 144 submerged electrodes addressing the 24 units of the plate sequentially through a multiplexing circuit for full spectrum (100 Hz to 200 kHz) sweeping and impedance measuring of the whole plate within 3 minutes. The AKITA Lid uses the tetrapolar configuration for impedance measurement, two or three pairs of electrodes were dipped into 3 reservoirs of each unit containing medium, acting as the current injection and voltage measurement electrodes. Figures 3a & 3d illustrate the two setups of electrodes addressing both sides of the membrane for TEER measurements. It is noted that the electrode characteristics play an important role in the signal quality. Hence, we performed several tests to ensure that the AKITA Lid can give reliable results. The tests include measurement sensitivity due to electrode placements and electrode dimensions and comparison with commercially available TEER measurement equipment.

**Figure 3.**
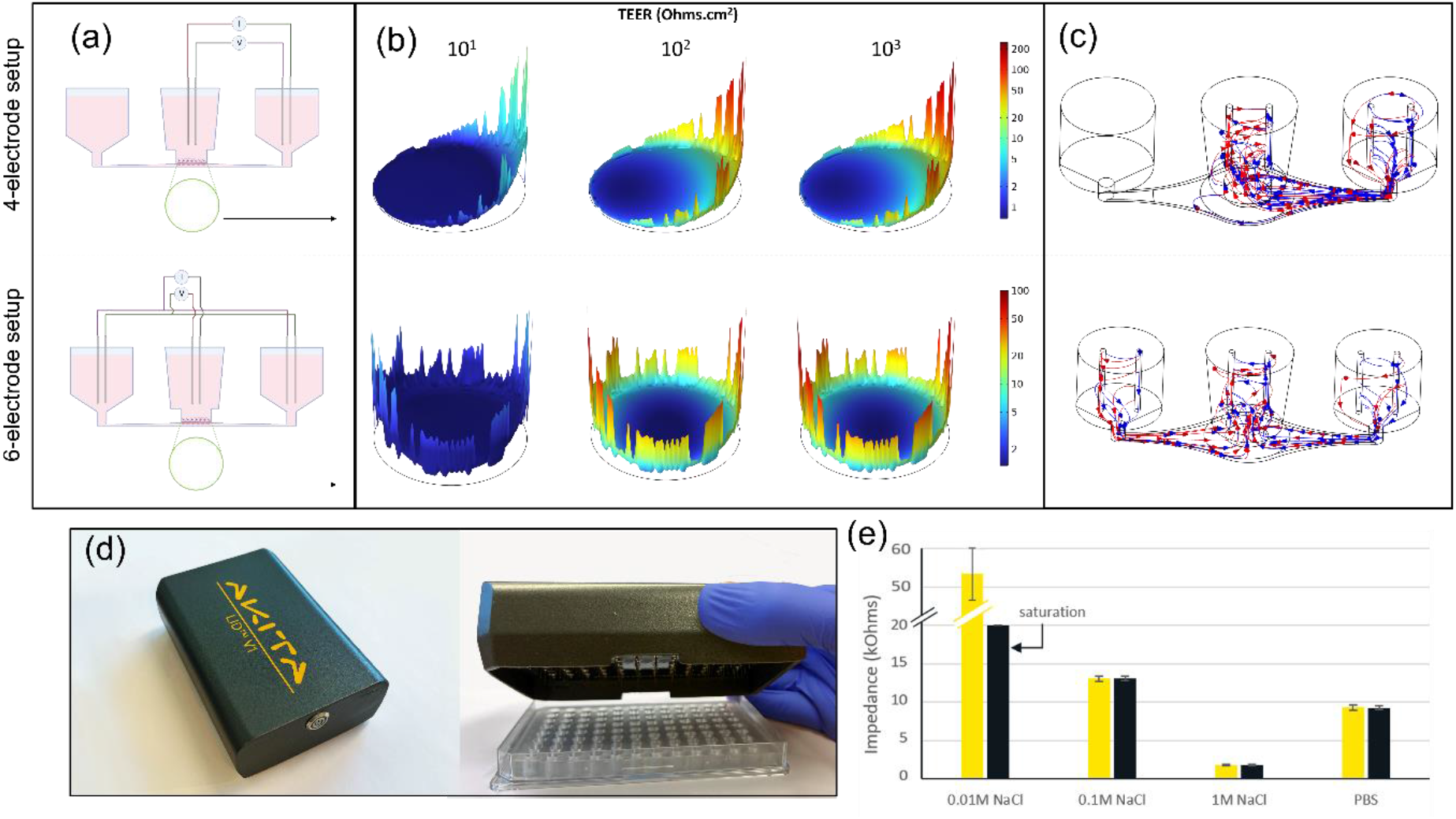
AKITA Lid TEER device. (a) A single cell culture unit with submerged chopstick-like electrodes. (b) Normalized planar sensitivity distribution (%) at 10 kHz of the membrane surface at different TEER values for 0.4 mm diameter gold electrodes. In both electrode setups, sensitivity peaks are observed on the edges of the cell surface and the sensitivity valleys are present at the center of the cell surfaces, but a uniform sensitivity is found over the barrier surface in 6-electrode setup at low TEER values (10 Ω.cm2). Note the logarithmic scale is different between the two electrode setups. (c) Current density streamline plot for the normal current (blue color) and the reciprocal current (red color). (d) The AKITA Lid is in operation with the AKITA Plate 96. (e) Impedance measurement made by the AKITA LID and the EVOM1 for solutions of varying conductivity: 1X PBS versus 0.01, 0.1, or 1M NaCl. Impedance is measured at 12.5 Hz for the EVOM and is the average of the impedance values between 500Hz and 30kHz for the AKITA Lid. N=4

The electrode’s geometry, arrangement, and microfluidic design have a great influence on the electric field during the TEER measurements ^33,34^. Over the area of interest, if there is a non-uniform electric field, only some portions of the cell layer contribute to the inferred TEER values, which do not represent the value of the whole cell layer. Planar sensitivity distribution on the cell layers on the AKITA Plate’s membrane was simulated with COMSOL using the reciprocal theorem ^35^. The sensitivity distribution was simulated at TEER values ranging from 10^1^ to 10^3^ Ω.cm^2^, covering the TEER range of most of the in vitro organ models ^36,37^. Figure 3b shows the normalized planar sensitivity (%) with height expression on the cell layer at 10 kHz frequency for a 4-electrode setup (Fig. 3a), with one pair of electrodes submerged in the open-top chamber, and the other pair inserted in the reservoir on the right side. Because of the lack of electrodes on the left reservoirs, electric current flow between the two electrode pairs tends to concentrate more on the right side (Fig. 3c), resulting in the lowest sensitivity observed on the left area of the membrane. Hence, we added two extension electrodes into the left reservoirs, merely a simple duplication of the other two electrodes in the right reservoir, forming a 6-electrode setup (Fig. 3d) to distribute the electric field across the membrane more evenly (Fig. 3f).

With 6 electrodes in total for the tetrapolar configuration, the simulation results showed a symmetrical distribution (Fig. 3e) of the sensitivity on the membrane and lower down the electrical resistance caused by the microfluidic channel. Furthermore, the 4-electrode TEER simulation results by Helm et al. have shown that the measurement sensitivity varies for different TEER values at different sweeping frequencies. For low-frequency sweeping, a uniform sensitivity is noticed on the cell layer with high TEER values. In contrast, at high-frequency sweeping, a more uniform sensitivity is observed on the cell layer at lower TEER values. Sensitivity is directly proportional to the TEER at higher frequencies whereas, at very low frequencies, sensitivity is inversely proportional to TEER ^38^.

Importantly, as observed in our simulation results, the non-uniformity still occurs even with the 6-electrode setup, and it was also mentioned elsewhere that the chopstick-like electrodes could not form a homogeneous electrical field along the cell layer ^39^. Both Zoio et al. and Yeste et al. simulated the sensitivity of the chopstick STX2 electrodes (World Precision Instruments) with 6.5mm and 12mm Transwell inserts, and both had non-uniformity of the sensitivity with “sensitivity valleys” ^33,40^. With AKITA Lid, two mitigation steps can be performed in the future to resolve the problem. First is to incorporate the electrodes into the AKITA Plate 96, to reduce the distance from the electrodes to the membrane and the distance between electrodes, providing a more uniform electric field distribution. The second is to widen the sweeping frequency range, to cope with the tissue barriers at low and high TEER values. Further simulation studies are required for optimizing the parameters such as electrode arrangement (distance between the electrodes), electrode shape, and electrode positioning (distance between the electrode tip and cell layer) for attaining a more uniform sensitivity at the cell layer.

Despite the drawbacks, the AKITA Lid illustrates a significant improvement compared to the conventional TEER measurement systems using manual single chopstick-like electrodes with low throughput and high variations ^33^. Furthermore, with a wide frequency range, AKITA Lid can perform measurements at much higher TEER values compared to conventional fixed-frequency TEER measurement systems.

To validate our AKITA Lid, we performed TEER measurements on the AKITA Plate 96 filled with buffers at different ion concentrations and compared it to the same measurements by EVOM1 (World Precision Instruments). To ensure measurement consistency, both devices should have the same electrode geometry ^41^. We thus customized an electrode connector to interface the EVOM1 with our electrode array, to fit with the AKITA Plate 96. Figure 3h shows the impedance measured for the 0.01M, 0.1M, and 1M NaCl, as well as 1X PBS. As expected, as the ions concentration increases, the impedance decreases. The AKITA LID measured an impedance more than four times higher for the 0.01M NaCl buffer compared to the 0.1M NaCl, and an impedance about 30 times higher for the 0.01M NaCl buffer compared to the 1M NaCl buffer. The EVOM and our AKITA LID have consistent impedance measurement results, except for the least ion-concentrated buffer: the EVOM was saturated at 20 kOhms impedance due to its fixed frequency AC current (12 Hz). Both devices show very low standard deviation as our electrode unit ensured a fixed positioning in the well.

### 3.4. Blood-brain barrier-on-chip

The application versatility of the AKITA Plate allows multiple BBB modeling opportunities. For proof-of-concept, we established both mouse and human BBB models for high molecular weight compound screening and reversible barrier opening assay for enhanced compound delivery to the brain. BBB-on-chip models were established with endothelial cells in the microchannel co-cultured with species-specific astrocytes and pericytes (Fig. 4a).

**Figure 4.**
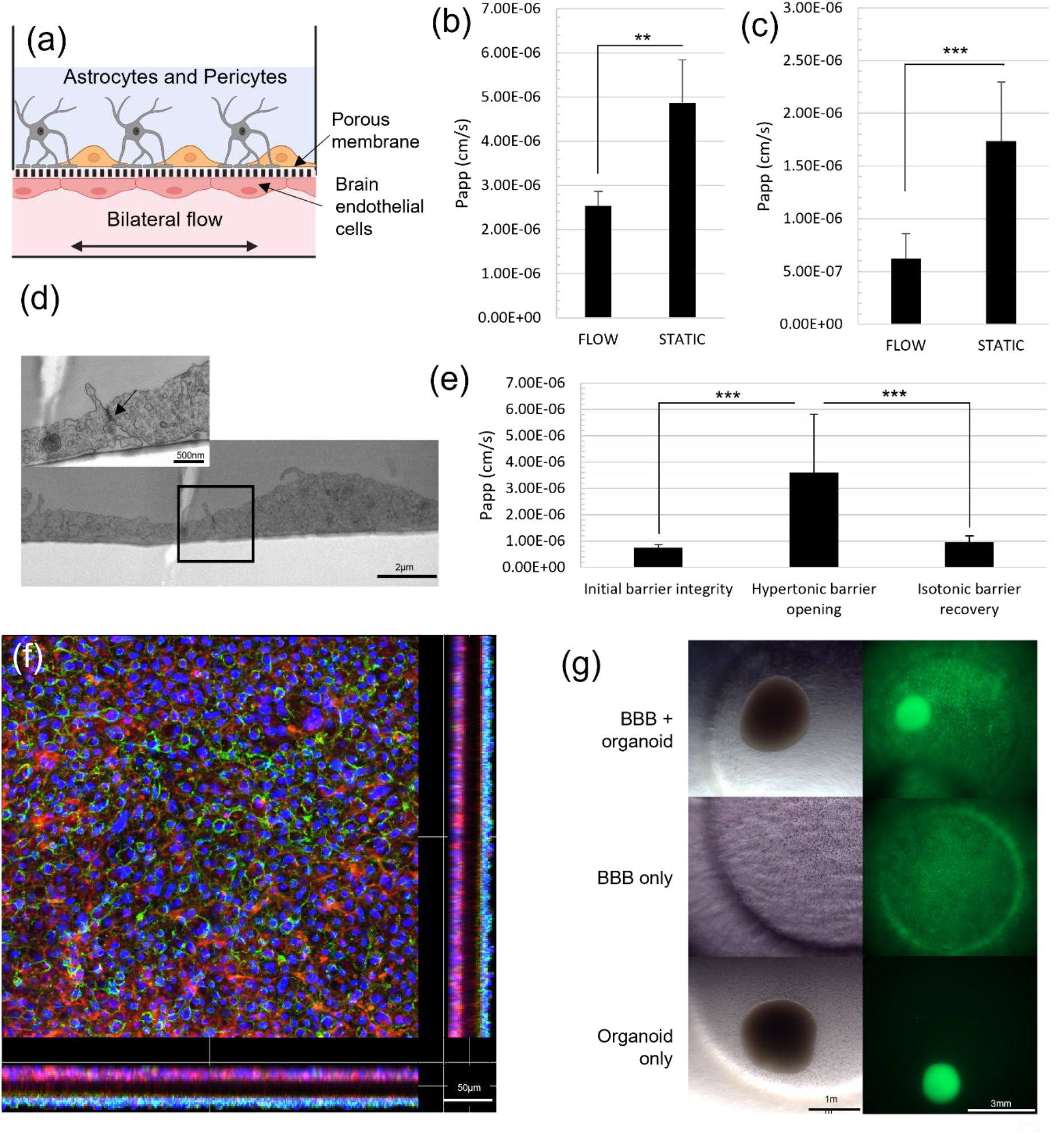
Blood-brain barrier-on-chip on AKITA Plate. (a) A scheme representing the BBB-on-chip model on AKITA® Plate with astrocytes and pericytes co-cultured with brain microvascular endothelial cells on opposing sides of the porous membrane. Both human (hBBB) and mouse (mBBB) models have tricultures of species-specific cells. (b) Apparent permeability (P_app_) of 70kDa dextran tracer for hBBB model cultured on AKITA Plate with and without dynamic flow. (c) P_app_ of 70kDa dextran tracer for mBBB model cultured on AKITA Plate with and without dynamic flow. (d) Transmission electron microscopy image from tight junction structure of the endothelial cells of the mBBB cultured on AKITA Plate. Tight junctions between the endothelial cells are marked with a black arrow (e) P_app_ of 70kDa dextran during the reversible barrier opening assay for mBBB model. (f) Confocal image of the mBBB cultured on AKITA® Plate showing co-culture monolayers on opposing sides of the porous membrane. Tight junction expression can be seen only in the endothelium side of the mBBB model. Blue DAPI, green CD31, red phalloidin. (g) Live imaging of BBB and organoid co-culture on AKITA® Plate, brightfield and Calcein AM signals shown. Data were analyzed using Two-way ANOVA and Tukey’s multiple comparison tests, **p<0,01, ***p<0,001, N=3

The endothelial cells were exposed to bilateral flow to ensure tight barrier formation. On day 5 of the experimentation, we assessed the permeability of the 70kDa dextran tracer through the triculture models of the human BBB (hBBB, Fig. 4b) and the mouse BBB (mBBB, Fig. 4c). As previously reported, inducing the flow in the microchannel improved the barrier tightness significantly when compared to hBBB or mBBB culture in a static environment ^42^. We also observed by electron microscopy that the endothelial cells are forming monolayers inside the microchannel (Fig. 4d). Additionally, the confocal microscopy analysis performed directly in the Akita plate showed the segregation of both cell populations at each side of the membrane. Indeed, the expression of the CD31 endothelial marker was specifically expressed at the bottom side of the membrane, while phalloidin stained the actin of both cell populations (Fig. 4f).

To enhance the delivery of drug compounds into the brain, transient opening of the BBB via the infusion of hypertonic solution can be performed both clinically ^43^ and in rodents ^44^. We aimed at reproducing this clinical protocol in our barrier model and optimized a reversible barrier opening assay using the hyperosmolar solution for BBB disruption. For this purpose, we established an mBBB with a coculture of mouse primary endothelial cells and mouse primary astrocytes. On day 7 of the experiment, the initial tightness of the barrier was verified to be close to values observed in vivo by using 70kDa dextran as a tracer (Fig. 4e) ^45^. After exposure to a hyperosmolar culture medium, we observed the transient opening of the barrier followed by recovery of the permeability scores after 1 hour back to the initial state (Fig. 4e).

Previously reported models have also shown successful cultures of BBB and midbrain organoids on transwell. The AKITA platform is compatible for organoid culture, hence we established a BBB-midbrain organoid co-culture model in a scalable throughput, with enhanced physiological flow conditions. First, we established a simple hBBB model with the monoculture of hCMEC/D3 cells in the microchannel and cultured the cells for seven days under dynamic conditions to allow barrier maturation. On the 7th day of the experiment, we added a midbrain organoid to the open-top chamber and monitored the viability of the coculture for an additional two days with calcein AM staining and live cell imaging (Fig. 4g). The viability of the organoid and the hBBB remained stable over the 48h co-culture, which provided a sufficient time window for the treatment with additional compounds.

### 3.5. Skin-on-chip

The structure of our full-thickness skin-on-chip model is displayed in Fig. 5a with a basal layer of dermal fibroblast covered by keratinocytes. Because high amounts of calcium have been shown to increase the maturation of keratinocytes, both high and low concentrations have been tested in our skin-on-chip model ^46^.

**Figure 5.**
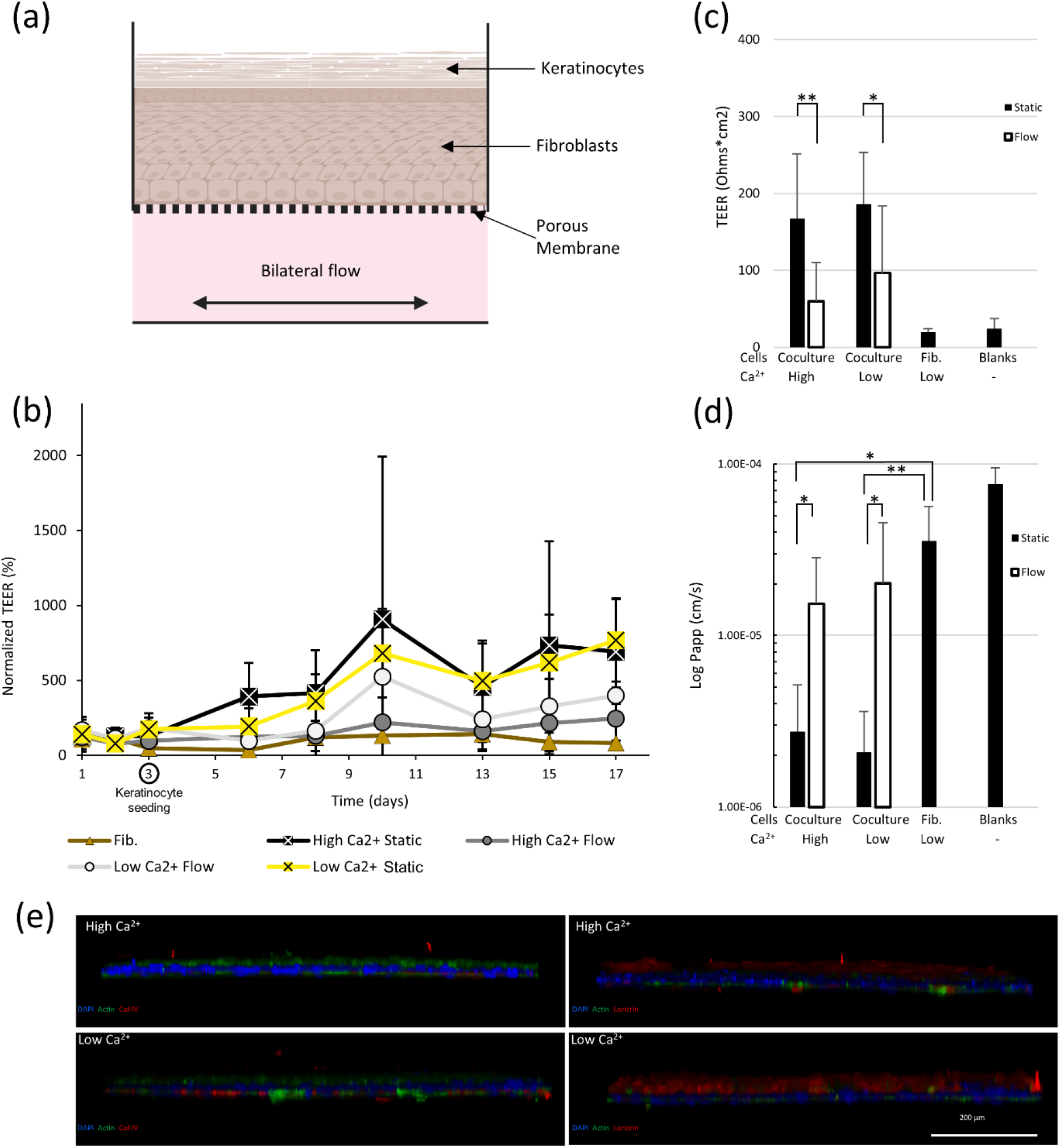
Skin-on-chip on AKITA Plate. (a) A scheme representing the skin-on-chip model in AKITA® Plate. Fibroblasts and keratinocytes are seeded on top of each other and cultured under the air-liquid interface. (b) TEER values for units measured during the model maturation up to 17 days, day 0 corresponds to fibroblast seeding. The units are cultured with or without dynamic flow, as well as with or without extra Ca^2+^ in the culture medium (High/Low). As a control fibroblast alone cultivated in static condition and low Ca^2+^ are displayed here (Fib.). (c) The TEER values acquired at tissue maturation, day 17 are plotted for statistical analysis. The no-cell control (blank) is available for comparison. (d) Apparent permeability values for the same skin-on-chip units measured with 0,4 kDa dextran. (e) Cross-sectional confocal images from stained skin-on-chip models cultured either with high or low Ca^2+^ medium. Stained with DAPI (blue), phalloidin (green), and collagen IV (left panel, red) or loricrin (right panel, red). N=3 for monocultures and N =12 for cocultures and no-cell controls (blanks) except N =6 for coculture units with low Ca^2+^ medium with 4 kDa permeability. Data were analyzed using two-way ANOVA tests and Tukey’s multiple comparison tests for dextran permeability and Kruskal-Wallis test together with the Dunn test for panel (c).

The dermal fibroblasts were cultivated for three days prior to the addition of keratinocytes. During the 17 days of tissue maturation, barrier integrity was monitored by TEER and showed a progressive tightness increase over time (Fig. 5b). After 14 days of co-culture required for keratinocyte maturation ^47,48^ (experiment day 17), barrier permeability was compared between TEER (Fig. 5c) and the diffusion of 0.4 kDa low molecular weight dextran from top to bottom (Fig. 5d). We noticed a high correlation between both TEER and dextran measurements (Fig. 5b-c). While both datasets showed a low impact of calcium concentrations on barrier permeability, an increased permeability under flow condition was observed (Fig 5. b-c). Those results indeed correlate with the poor physiological relevance of having flow in skin tissue; the addition of an endothelial cell layer may be required to elicit improved flow - and physiological relevance - outcomes.

Finally, the stratification of the skin tissue was observed by confocal microscopy in the Akita plate. Keratinocyte cornification was monitored with Loricrin ^49^ while Collagen IV labels the collagen basal membrane ^50^ (Fig. 5e). Both stains show higher tissue stratification in the low calcium concentration which contrasts with previous studies ^46^. In our system, the air-liquid interface only allows the delivery of culture media to the basal fibroblasts, which may impair the latter. Such observation confirms that our skin model may be used to study impairment in skin stratification, barrier integrity, and compound absorption.

### 3.6. Lung-on-chip

The application capacity of a high-throughput platform for lung modeling was optimized with the co-culture of Adenocarcinoma human alveolar basal epithelial A549 cells and human umbilical vascular endothelial cells (HUVEC) as depicted in Figure 6a. Confocal microscopy analysis of a day 10 epithelium cultivated in an air-liquid interface shows the expression of the maturation marker mucus protein MUC5AC on the apical side of the epithelium (Fig. 6b).

**Figure 6.**
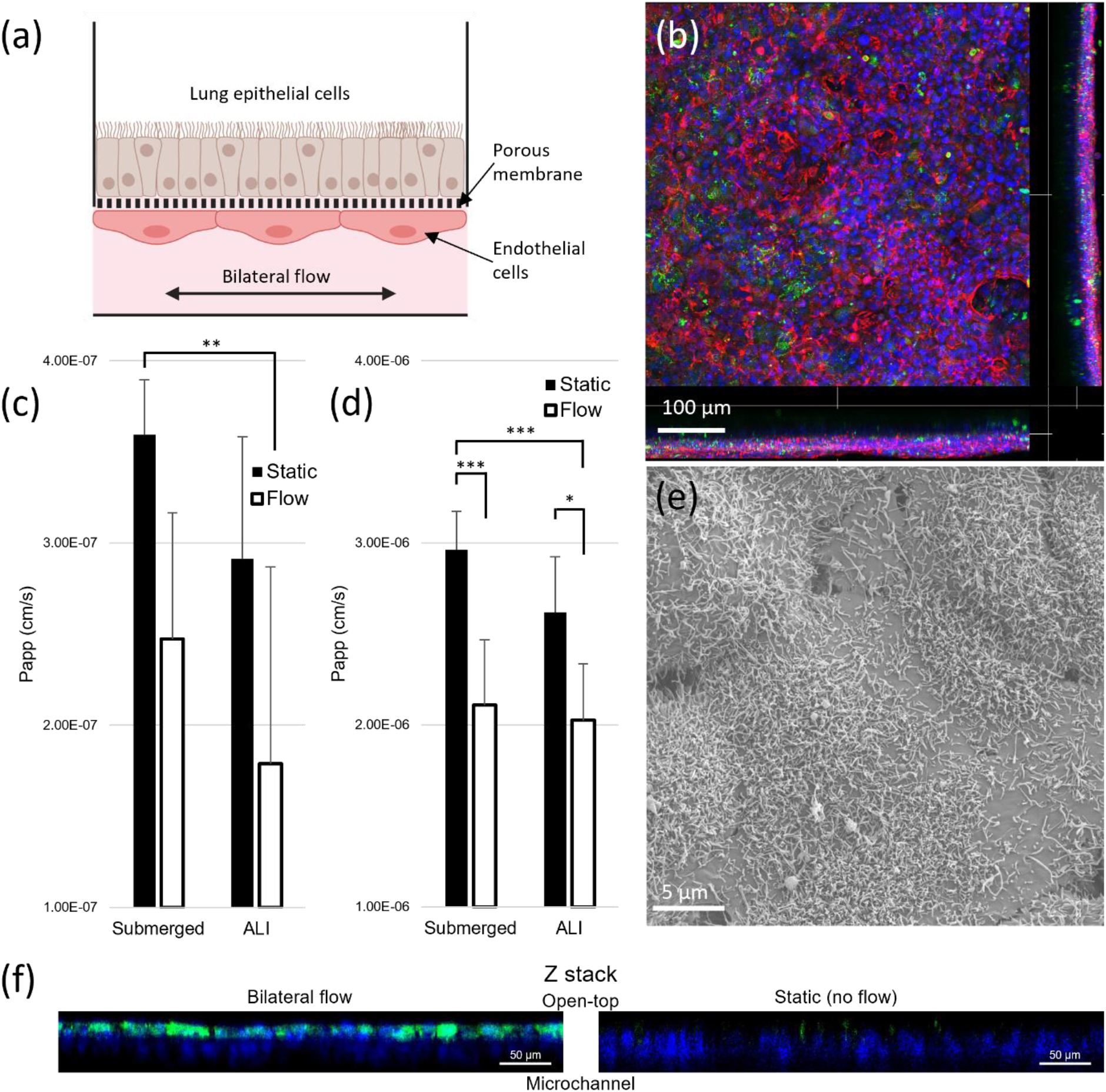
Lung-on-chip on AKITA Plate. (a) Scheme representing the lung-on-chip model in AKITA® Plate. Lung epithelial cells (A549) and endothelial cells (HUVEC) are seeded on the other side of the porous membrane. (b) Confocal image of the A549 on the lung-on-chip barrier. Blue DAPI, green MUC5AC, red phalloidin. (c) Apparent permeability values (Papp) for lung-on-chip units cultured with or without dynamic flow, and with or without an air-liquid interface (ALI) measured with 70 kDa dextran tracer. (d) Apparent permeability values (Papp) for lung-on-chip units cultured with or without dynamic flow, and with or without an air-liquid interface (ALI) measured with 4 kDa dextran tracer. (e) SEM image from microvilli structure on top of the epithelial layer cultured under the air-liquid interface in AKITA Plate. (f) Cross-sectional confocal images of lung-on-chip barriers cultured either in dynamic flow or static environment. Stained with DAPI (blue) and MUC5AC (green). Data were analyzed using Two-way ANOVA tests and Tukey’s multiple comparison tests. N=6.

Additionally, we show already after 2 days of air-liquid interface culture that the combination of both flow and air-liquid interface improve barrier tightness observed with the 70 kDa dextran tracer (Fig. 6c). Surprisingly, for the 4 kDa lower molecular size tracer, both flow and air-liquid interface increased the tightness of the model even alone (Fig. 6d). Overall, dynamic flow seems to have a higher effect on barrier tightness, as reported previously ^51^. With the 4kDa dextran tracer, our model appears to achieve higher tightness compared to previous studies which obtained 8,44E-6 cm/s ^52^, 1.70E-5 cm/s ^53^. While with the 70kDa tracer our models show comparable permeability compared to literature 3,8E-6 cm/s ^53^ and 6,98E-8 cm/s ^54^. In contrast with Chang et al. ^55^, the air-liquid interface had a limited impact on the tightness of our system, likely due to the use of different cell lines in the studies.

The A549 are type II cells that exhibited the formation of microvilli on their apical side observed by electron microscopy (Fig. 6e) ^56,57^. Together with the apical expression of the mucus protein MUC5AC, (Fig. 6f), this further confirmed the polarization of the lung epithelium and demonstrated its functionality. As can be seen from Figure 6f, dynamic flow also increased mucus production compared to the static culture. This model may be further used for the high throughput assessment of drug compound absorption, the test of anti-inflammatory compounds, or the effect of pro-inflammatory stimuli and pollutants on lung-on-chip barrier integrity.

## 4. Conclusions

We present a versatile and scalable OOC platform, providing a low to high-throughput reproducible solution for in vitro tissue barrier modeling with pumpless flow control. It may be used to study the pharmacodynamics of drug compounds crossing biological barriers, and barrier integrity via rapid label-free TEER measurements, of high-content screening with confocal and standard fluorescent microscopy. The current laboratory equipment such as microscopes, automated liquid handling robots, high-content screening systems, etc. can adopt the AKITA Plates into their workflows seamlessly thanks to the standard design and simplicity of the plate handling.

